# Integrating Phylogenies with Chronology to Assemble the Tree of Life

**DOI:** 10.1101/2024.07.17.603989

**Authors:** Jose Barba-Montoya, Jack M Craig, Sudhir Kumar

## Abstract

Reconstructing the global Tree of Life necessitates computational approaches to integrate numerous molecular phylogenies with limited species overlap into a comprehensive supertree. Our survey of published literature shows that individual phylogenies are frequently restricted to specific taxonomic groups due to the expertise of investigators and molecular evolutionary considerations, resulting in any given species present in a minuscule fraction of phylogenies. We present a novel approach, called the chronological supertree algorithm (Chrono-STA), that can build a supertree of species from such data by using node ages in published molecular phylogenies scaled to time. Chrono-STA builds a supertree of organisms by integrating chronological data from molecular timetrees. It fundamentally differs from existing approaches that generate consensus phylogenies from gene trees with missing taxa, as Chrono-STA does not impute nodal distances, use a guide tree as a backbone, or reduce phylogenies to quartets. Analyses of simulated and empirical datasets show that Chrono-STA can combine taxonomically restricted timetrees with extremely limited species overlap. For such data, approaches that impute missing distances or assemble phylogenetic quartets did not perform well. We conclude that integrating phylogenies via temporal dimension enhances the accuracy of reconstructed supertrees that are also scaled to time.

## 1 Introduction

Reconstructing the history of life on Earth is foundational to studying evolution and biodiversity, which is pursued by many taxonomists, systematists, and evolutionary biologists. Molecular phylogenetics has been a key tool to infer the evolutionary relationships of organisms (Hedges and Kumar, 2009; Yang and Rannala, 2012). Occasionally, large phylogenies are constructed by extensive sampling of species from major groups like birds, squamates, mammals, and fishes (Jetz *et al*., 2012; Upham *et al*., 2019; Tonini *et al*., 2016; Hughes *et al*., 2018; Álvarez-Carretero *et al*., 2022) Yet, much more commonly, published phylogenies are the work of taxon specialists who focus on individual families or genera due to their organismal expertise. Furthermore, even considering the increased accessibility of genetic data and improvements in computational power, technical impediments still stand in the way of building large-scale phylogenies. For example, while certain genetic loci contain valuable phylogenetic signals in some taxa, they may be largely invariant or actively misleading in others (Gonçalves *et al*., 2019). Moreover, teasing apart orthologous from paralogous sequences can be challenging, especially among increasingly distantly related taxa (Koonin, 2005; Altenhoff *et al*., 2019). Similarly, the best models to capture the processes of sequence evolution in one clade may be inappropriate for another (Lopez *et al*., 2002; Author and Fitch, 1971; Kumar *et al*., 2005). Therefore, many small and large taxonomically restricted phylogenies have been published (Hedges *et al*., 2015; Kumar *et al*., 2022).

Instead, this collection of phylogenies, with so little taxonomic overlap, could be combined by using the species divergence times because all these trees were scaled to absolute time (timetrees) (**Figure 2**; Chrono-STA). In this case, divergence times were used to merge species into a supertree by first connecting the most closely related species, i.e., species pairs with the smallest divergence time and then repeating this step iteratively. Therefore, incorporating chronological information can mitigate the extremely limited and uneven overlap observed in the collections of empirical timetrees.

Hedges et al. (2015) have previously described a hierarchical average lineage (HAL) clustering based on divergence times to resolve polytomies in the NCBI backbone taxonomy, followed by localized branch swapping to make evolutionary relationships maximally consistent with the topologies. While this approach could lead to the assembly of a supertree of more than 148,000 species from published phylogenies (Hedges *et al*., 2015; Kumar *et al*., 2022), HAL is limited by its requirement of a phylogenetic backbone that creates many additional polytomies in cases where the sample of input trees conflict with the backbone and one another, which we found it to be a common problem. Instead, there is a need for a principled algorithm for the *de novo* reconstruction of a supertree from the collection of timetrees.

Here, we present a chrono-supertree algorithm (Chrono-STA) that does not require a phylogenetic backbone to build a supertree from a collection of timetrees. It pairs species using all the input timetrees analyzed in parallel independently. Chrono-STA does not need to construct or reduce a global matrix of divergence times between species like HAL does (see *Material and Methods*, section 2.1). Also, Chrono-STA does not impute missing nodal distances between taxa, such as those in some methods (Morel *et al*., 2022; Vachaspati and Warnow, 2015).

In the following, we first present the concept and implementation of Chrono-STA, then demonstrate its usage by analyzing three computer-simulated datasets and one empirical dataset. In these examples, timetrees have very few common species to mimic the patterns observed in the corpus of published timetrees (**Figure 1**). We also applied some gene tree conciliation approaches to these datasets for assessing the relative performance of methods based on different approaches to combining phylogenies with partial overlaps.

**Figure 1.**
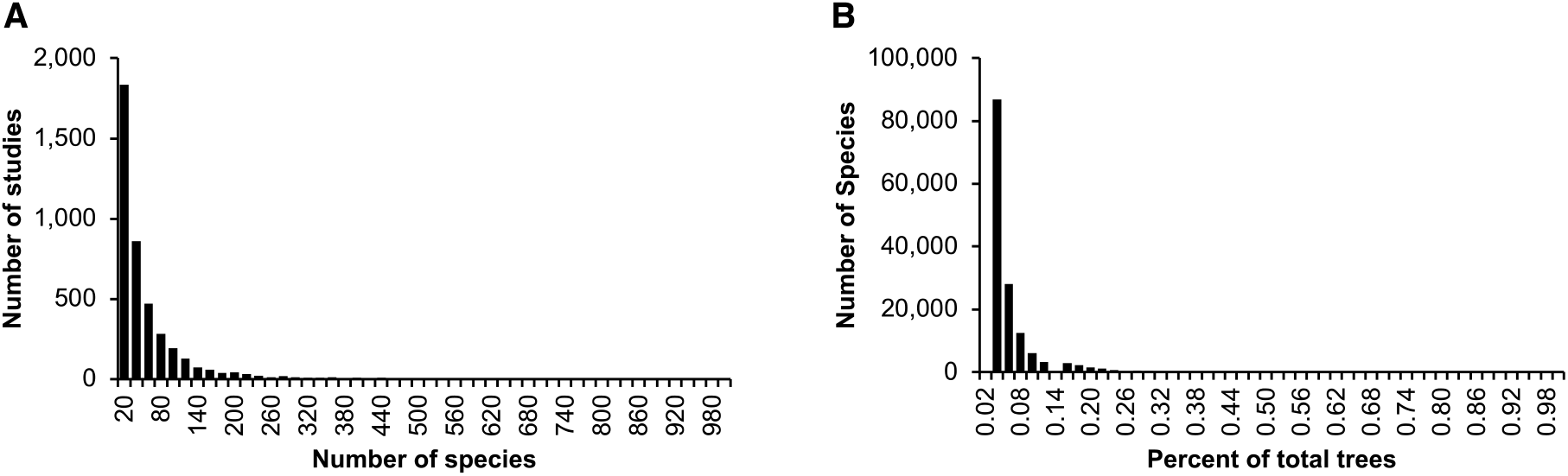
Summary characteristics of published timetrees curated for the TimeTree database (Kumar *et al*., 2022). Distributions are shown for the (**A**) number of species in phylogenies, (**B**) count of phylogenies in which a species occurs, as a percentage of the total number of trees. These statistics were derived from 4,185 phylogenies published in the last five decades. Only species counts up to 1,000 (**A**) and percentages up to 1% (**B**) are shown. Fundamental properties of published species trees can be gleaned from the collection of more than 4,000 phylogenies curated for the TimeTree database (Kumar *et al*., 2022). Across the whole collection, phylogenies contained a median of 25 species each (**Figure 1A**), each found in a median of just one timetree (0.02% of the sample) (**Figure 1B**). Consequently, the average number of species common between any two phylogenies is less than 1. Uniting a collection of phylogenies with such limited taxonomic overlap is not the intent of existing methods to combine phylogenies. The difficulty is illustrated through an example in **Figure 2** for a collection of five timetrees (trees 1 to 5) derived from a modal tree of seven distinct species (**Figure 2**, model tree; species A to G). We applied existing methods for reconciling gene trees, which either impute missing nodal distances between species (e.g., Asteroid (Morel et al., 2022) and ASTRID (Vachaspati and Warnow, 2015)) or reconcile quartets (ASTRAL-III; Zhang et al., 2018). None of them could recover the correct tree for this example collection (**Figure 2**). This is because of the sparsity of the species overlaps between trees.

**Figure 2.**
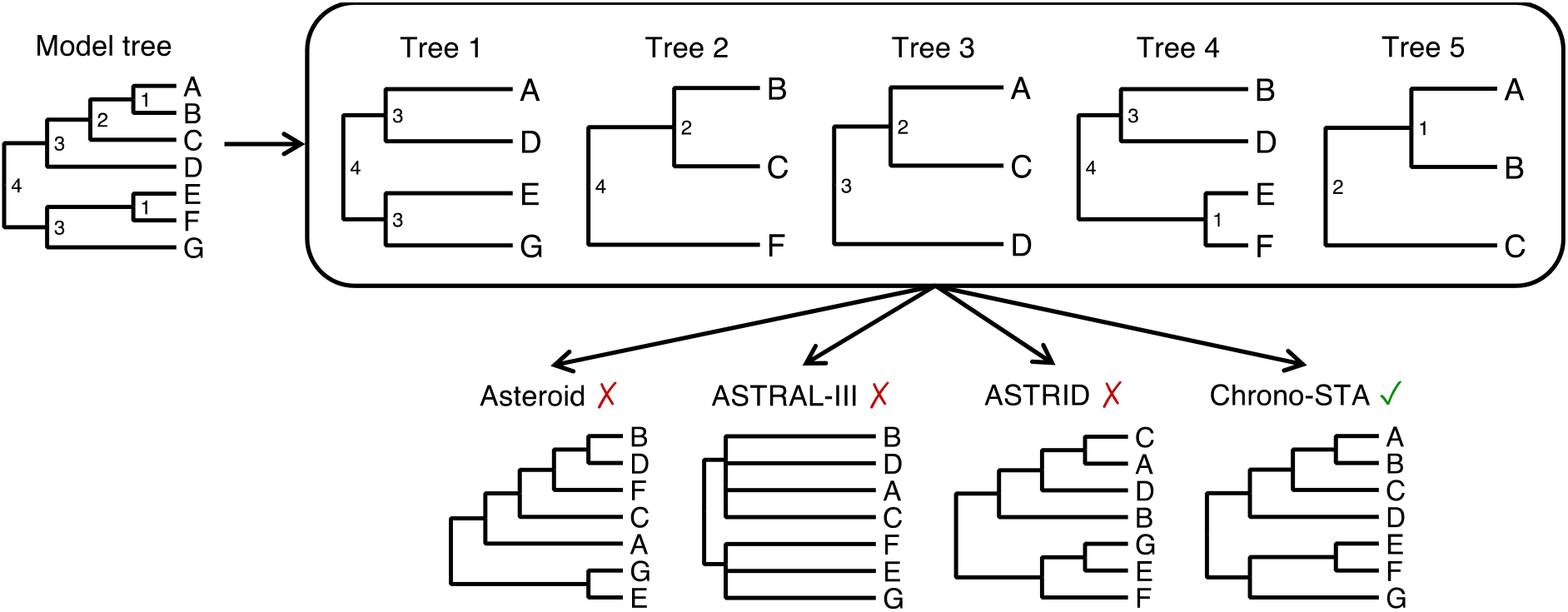
Five timetrees with partial species overlaps, derived from a model tree by randomly sampling taxa (node ages are shown to the right of each node). Supertrees were produced by gene tree reconciliation methods that can handle missing data: Asteroid, Astral-III, and ASTRID. The new Chrono-STA approach was able to produce the correct supertree using divergence times.

## 2 Material and Methods

### 2.1 A novel chronological supertree approach (Chrono-STA)

In Chrono-STA, pairs of taxa with the least divergence time are clustered iteratively. This requires finding a pair of taxa with the smallest divergence time in all the timetrees in every step. If a pair of taxa is found in multiple timetrees, then the mean across all relevant timetrees is used to derive a summary estimate. Upon the identification of taxa with the smallest divergence time (say, *a* and *b*), they are clustered and represented by a unique label (say, *z*). Then, tips with labels *a* or *b* are replaced by their group name (*z*) in all the constituent timetrees. Now, the number of unique taxa in the collection of timetrees has decreased by one, whereas the global timetree consists of two taxa along with their divergence time.

Next, a pair of taxa with the smallest divergence time is identified from the new collection of timetrees following the procedure above. Notably, the calculation of mean divergence time for some pairs of taxa in some timetrees will require special attention. If a timetree has multiple tips with the same name (e.g., due to the grouping of *a* and *b*), we first obtain timetree-specific mean divergence times for the taxa pair of interest (e.g., *z* and *c*). Then, the taxa pair with the least divergence time is identified. Ties can be broken arbitrarily. This clustering and name replacement process is repeated until no pairs of taxa are left with a divergence time.`

At this point, we build a supertree by connecting all the groups based on taxon pairings made. If the species overlap fails to connect all the species into a single timetree, multiple sub-global timetrees may result. The final timetree may not be ultrametric because individual divergence times are estimated with variance and may suffer from systematic biases (Barba-Montoya *et al*., 2017; Hedges *et al*., 2018), so we subject the supertree to time-smoothing by using non-negative least squares.

For simplicity, we implemented the above approach in an algorithm that works with matrices of pairwise distances for each timetree (**Supplementary Figure 1**): (A) assemble a collection of timetrees, (B) compute a time distance matrix between taxa independently for each timetree, (C) compute averaged super matrix, (D) identify the smallest divergence time, select the taxa pair, (E) propagate the pairing to all the individual time matrices by replacing the paired taxa name by the new name and calculate pairwise divergence times between the new taxon and all the other taxa, repeat steps C-E until there are no more taxa pairs with divergence times, (F) generate a list of clusters and pairwise distance times (G) generate a complete pairwise time distances matrix including previously missing pairs, (H) connect all the clusters and pairwise distance times into a supertree, and make supertree ultrametric. This approach can also be applied to combine trees from (I) partially overlapped multi-sequence alignments (MSAs) by (J) inferring an ML tree for each MSA and dating each ML tree. Then, the constituent timetrees are combined following the outlined procedure.

### 2.2 Tested methods

Using simulated and empirical data, we compared the performance of Chrono-STA and four other supertree construction methods. Chrono-STA requires no fine-tuning parameters for analysis except for the collection of supertrees. For ASTRID (Vachaspati and Warnow, 2015), the FastME analysis was conducted with both nearest neighbor interchange (NNI) and subtree-pruning-and-grafting SPR moves (-s option), and (-u) to use UPGMA completion. For ASTRAL-III (Zhang *et al*., 2018), a heuristic search was conducted. Branches on the supertree were scored using the posterior probability for the main resolution (-t 3). The lambda parameter for the Yule (Yule, 1925) prior, used branch lengths and posterior probabilities (-c) calculations, was set to 0.5. For Asteroid (Morel *et al*., 2022), a heuristic search was conducted to find the supertree with the lowest global induced length. Asteroid begins with a specified supertree topology and utilizes a tree search strategy, incorporating SPR moves to optimize the score. We used 20 randomly generated starting trees (-r 20). The supertree topology was iteratively optimized by an adapted FastME (Lefort *et al*., 2015) tree search algorithm to the global induced length score.

#### 2.2.1 Quantifying and comparing performance

The performance of the methods for constructing supertrees was assessed by calculating Robinson-Foulds (RF) distances between the inferred and reference tree (Robinson and Foulds, 1981). This calculation is performed using the R function MultiRF (Revell, 2012). The normalized Robinson-Foulds (nRF) distance estimates the topological error in phylogeny reconstruction. It is calculated as nRF = RF / (2(*n*-3)), where *n* is the number of species. The model timetree was the reference tree for simulated datasets, whereas the timetree published in the original study was assumed to be the reference tree in the analysis of empirical data.

Chrono-STA also produces node ages in the inferred supertree, compared with the times in the reference tree. Because the topologies of inferred and reference phylogenies were not identical, we compared the node times in the reference tree with that of the most recent common ancestor (MRCA) in Chrono-STA timetree of the set of taxa that descended from that node in the reference tree. The *slope* and coefficient of determination (*R*^2^) for the linear regression through the origin were computed for the comparison of the inferred supertree and the reference tree. Furthermore, the difference between the estimated MRCA node times and reference tree node times was computed. The difference was divided by the reference tree node time and multiplied by 100 to generate a percent time error (ΔTE).

### 2.3 Datasets

#### 2.3.1 Simulated datasets

To assess the performance of Chrono-STA in constructing supertrees from timetrees with extremely low species overlap, three small collections (C1-C3) of six timetrees (T1-T6) were generated (**Figure 3**). Each timetree was derived from an alignment of 51 species from the collection of sequence alignments utilized previously by Tamura et al. (2012). They generated alignments using SeqGen (Rambaut and Grassly, 1997) under the HKY substitution model (Hasegawa *et al*., 1985) and heterogeneous sets of evolutionary parameters, including sequence lengths (258 to 9353 sites), evolutionary rates (ranging from 1.35 to 2.60 substitutions per site per billion years), G+C-content bias (G+C contents ranging from 39% to 82%), and transition/ transversion rate bias (transition/transversion ratio, ranging from 1.9 to 6.0. We selected six nucleotide gene alignments (A1-A6; ranging from 2,174 to 3,100 sites) simulated with autocorrelated rate variation among lineage, such that the rate of a descendant branch was drawn from a lognormal distribution centered around the mean rate of the ancestral branch; an autocorrelation parameter *v*=1 was used (Kishino *et al*., 2001). Their original datasets contained 446 species, but we sampled 51 species, as in Barba-Montoya et al. (2023), for practicality (**Figure 3**). During species down-sampling, an outgroup as well as at least one of the ingroup root taxa selected to ensure that timetrees could be produced from sequence alignments. The simulated datasets and model timetree are available at https://github.com/josebarbamontoya/chrono-sta.

**Figure 3.**
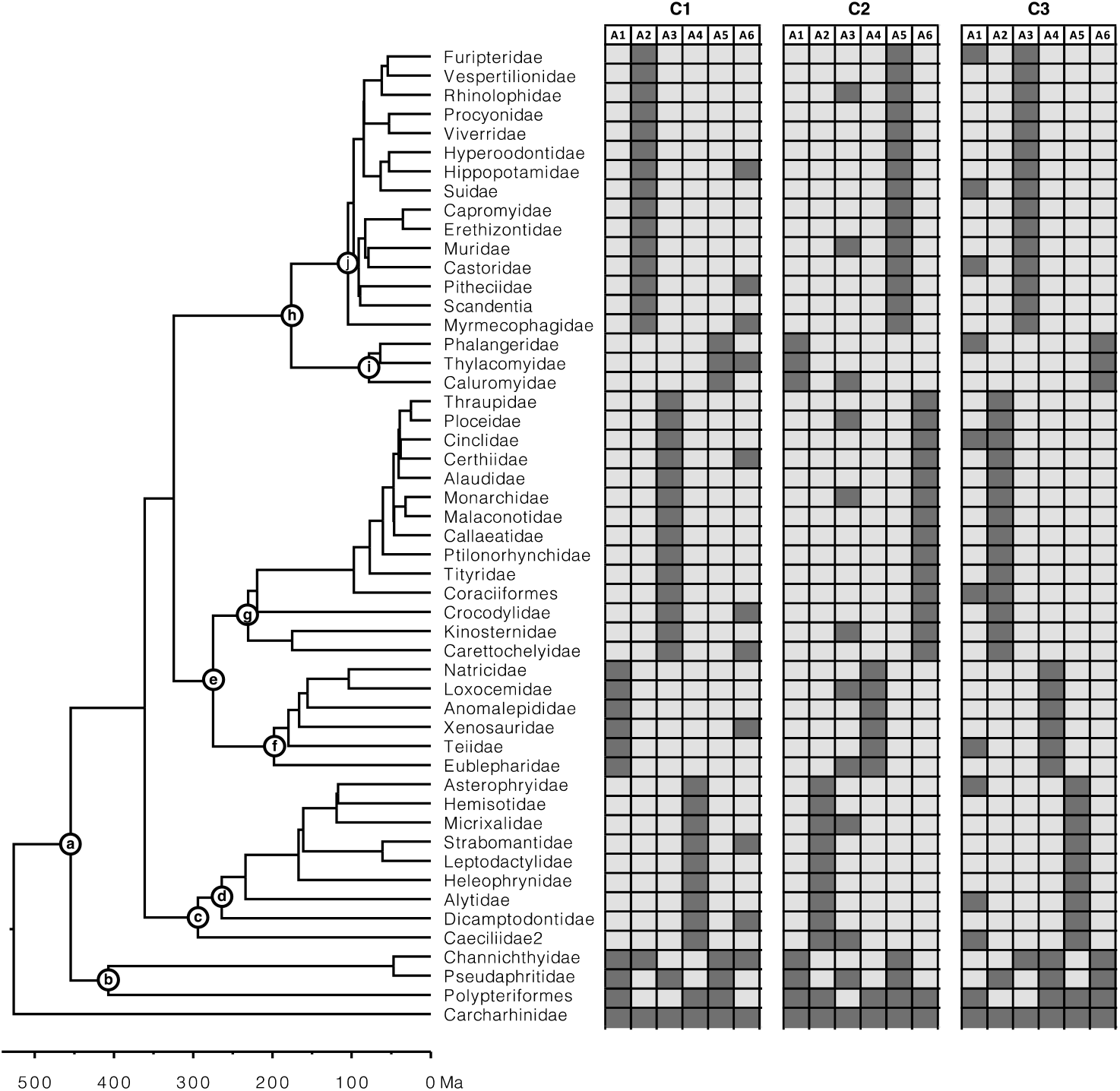
Presence-absence representation of the three simulated collections (C1-C3) of six gene alignments (A1-A6) each, with extremely low overlap of species, generated to construct supertrees from constituent trees. Each collection consists of five constituent timetrees along a backbone timetree. The calibrations (nodes a-j) used to construct the constituent timetrees are represented on the model timetree (reference tree). The rooting outgroup (Carcharhinidae) was excluded from the analysis.

Constituent timetrees were inferred using gene alignments for every collection (CI-C3). First, an ML tree was inferred from each gene alignment using the HKY+Γ5 model in IQ-TREE (Nguyen *et al*., 2015). Then, each ML tree was dated using RelTime (Tamura *et al*., 2012) in MEGA-CC (Kumar *et al*., 2012; Tamura *et al*., 2021). Each timetree was computed using a set of 10 calibrations, each assigning a uniform distribution: (a) U(453, 457 Ma), (b) U(405, 409 Ma), (c) U(292, 296 Ma), (d) U(262, 266 Ma), (e) U(273, 277 Ma), (f) U(196, 200 Ma), (g) U(229, 233 Ma), (h) U(174, 178 Ma), (i) U(76, 80 Ma), and (j) U(103, 107 Ma). The rooting outgroup (Carcharhinidae) was excluded from the analysis because RelTime analysis does not produce estimates in the outgroup (Tamura et al., 2018; 2012). Each collection contained five taxon-restricted timetrees with limited overlap and one timetree with one or more species per major group. These timetrees were derived from 51-species alignments by a realistic process that ensured that every collection’s individual timetree (T1-T6) contained phylogenetic errors and produced node ages with variance, as would be the case in real studies. The timetrees produced were missing an average of 78% of species, with a range of 67% to 88%, and had a limited species overlap.

#### 2.3.2 Empirical dataset

To assess the performance of methods in constructing supertrees from limited overlapping timetrees, we used the mammal timetree of Álvarez-Carretero et al. (2022) that consisted of 4,705 species across 14 constituent timetrees, including a backbone timetree (**Figure 4**). This collection of timetrees was combined using the supertree construction method with parameters set as described above. The 4,705 mammal species timetree and the 14 constituent timetrees are available at https://github.com/josebarbamontoya/chrono-sta.

**Figure 4.**
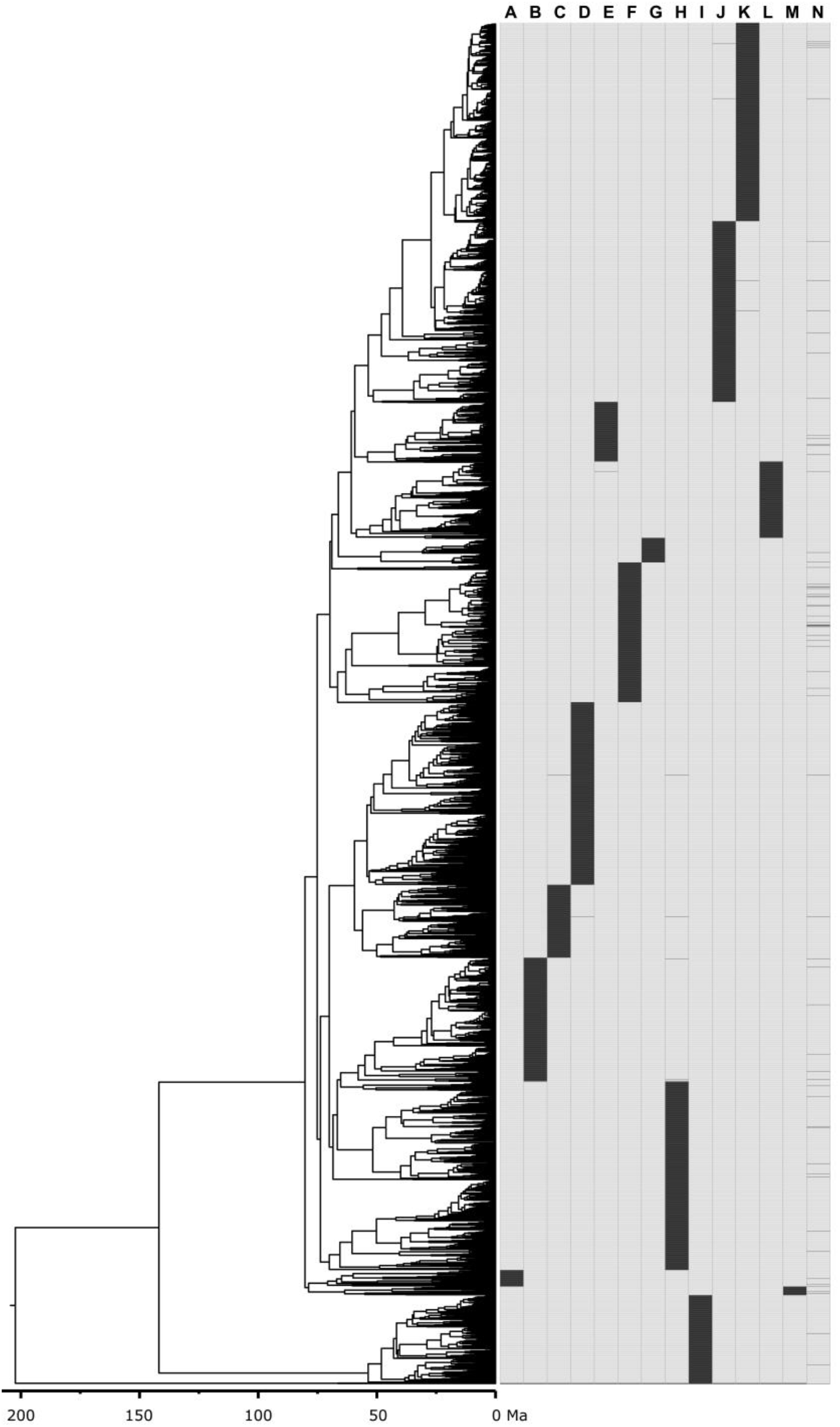
Mammal timetree consisting of 4,705 species across 14 constituent timetrees including a backbone timetree (Álvarez-Carretero et al., 2022). Presence-absence representation of 14 constituent timetrees: (A) Afrotheria, (B) Artiodactyla, (C) Chiroptera_subt1, (D) Chiroptera_subt2, (E) Ctenohystrica, (F) Euarchonta, (G) Lagomorpha, (H) Laurasiatheria_therest, (I) Marsupialia, (J) Rodentia_therest_subt1, (K) Rodentia_therest_subt2, (L) Sciuridae_and_related, (M) Xenarthra, (N) 00_main_tree_T2-updated-geochronolog (backbone timetree).

## 3 Results

### 3.1 Accuracy of supertrees constructed from constituent timetrees

We first assessed the performance of Chrono-STA for the simulated data (**Figure 3**). Five of the six timetrees in each of the three collections had excellent taxonomic coverage within clades, but only a limited overlap with other timetrees. 73% species occur in just one of five trees, while only one species is common to all the timetrees. This design mimics empirical phylogenies which often specialize on given clades. Individual phylogenies in each collection differ in topology and times, because every timetree was inferred independently from a simulated multispecies alignment, as described in the *Material and Methods* (section 2.3.1).

On average, Chrono-STA produced a supertree whose phylogeny agreed 90% with the reference tree, i.e., nRF = 0.1, from all three collections of timetrees (**Figure 5**). Therefore, Chrono-STA can work well for datasets with limited overlaps among major groups of taxa. ASTRAL-III achieved an average nRF of 0.2, which was twice as inaccurate as Chrono-STA. Other methods performed worse, with an average nRF of 0.42 for ASTRID and 0.54 for Asteroid. Overall, these results suggest that the inclusion of chronology while combining phylogenies can produce higher accuracies when species overlaps are limited. The incorporation of the time dimension is a fundamental unifying factor, which other methods do not use as effectively as Chrono-STA.

**Figure 5.**
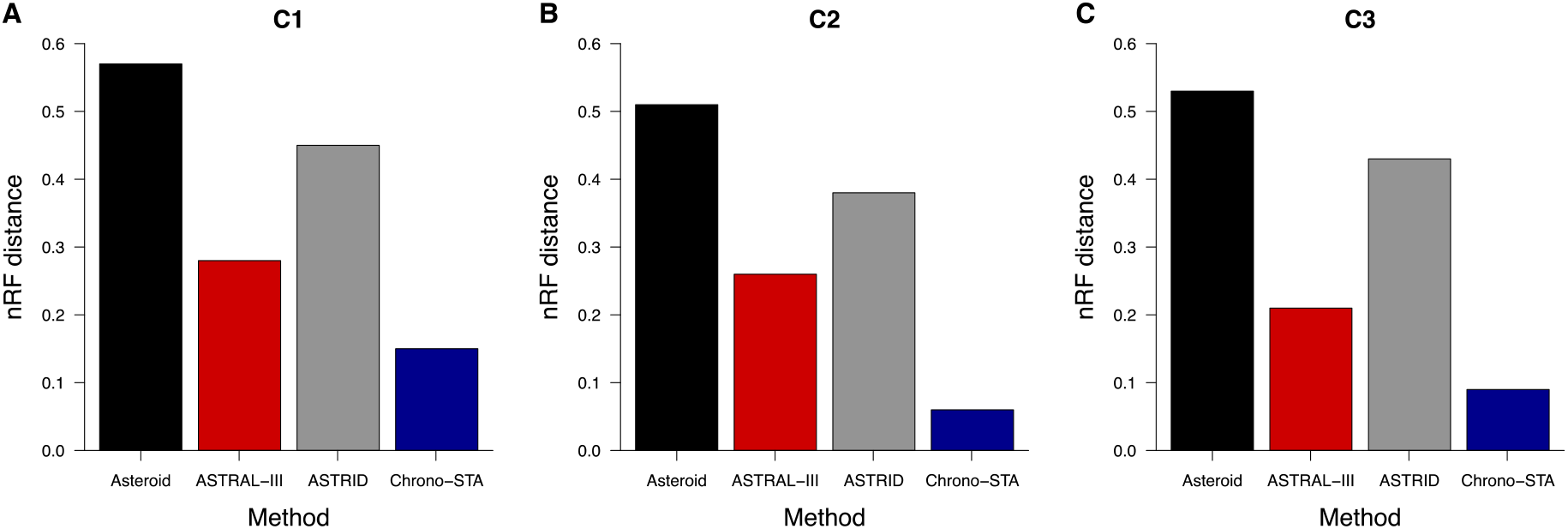
nRF distances between the model timetree (reference tree) and the generated supertree across collections (**A**) C1, (**B**) C2, and (**C**) C3 for Asteroid (black), ASTRAL-III (red), ASTRID (grey), and Chrono-STA (blue).

Chrono-STA produces divergence times along with the phylogeny. So, we compared the time estimates from the inferred Chrono-STA supertree with those in the reference tree. We used the Chrono-STA node times for the MRCA of all the sets of taxa in the reference tree. Chrono-STA generated node times highly consistent with those of the reference tree (**Figures 6A-C**), with *slope* and *R*^2^ values approaching 1.0 across all datasets. We also quantified the accuracy of Chrono-STA by computing the difference between the estimated MRCA node times and the true node times. The median ΔTE was low for the three datasets (**Figure 6D**), at –9% for C1, –1% for C2, and –0.5% for C3.

**Figure 6.**
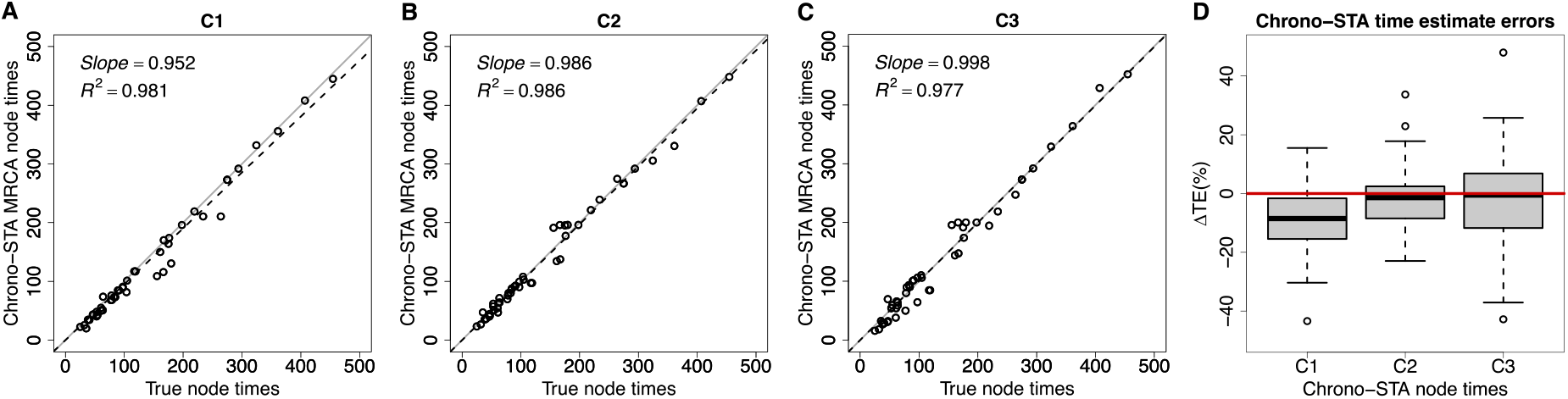
(**A-C**) Comparison of time estimates obtained by using Chrono-STA with the true node times for collections C1-C3. The *slope* and coefficient of determination (*R*^2^) for the linear regression through the origin are shown. The black dashed lines represent the best-fit linear regression through the origin. The solid gray line represents equality between estimates. (**D**) Distributions of the differences between Chrono-STA node times and true node times (ΔTEs). The black horizontal line represents the median value. For Chrono-STA, we used the estimated node times for the MRCA of all the sets of taxa in the model timetree (reference tree).

We validate these trends observed in simulated data by analyzing the large empirical dataset of Álvarez-Carretero et al. (2022), containing 4,705 mammal species across 14 taxonomically restricted timetrees (**Figure 4**). Chrono-STA assembled these timetrees into a supertree that was identical in topology to that published by Álvarez-Carretero et al. (2022) (**Supplementary Figure 2**), except for a single internal branch which shifted to its sister clade, indicated by a red and a black asterisk in **Figure 7A**. The nRF for Chrono-STA was 0.0002 (**Figure 7B**). ASTRAL-III performed the second best (**Figure 7B**), generating a supertree with 96 topological differences from the reference tree (**Supplementary Figure 3**), which is almost 200-times worse than Chrono-STA (nRF = 0.02). No other tested method performed well (**Figure 7B**). the ASTRID supertree had 430 differences from the reference tree (**Supplementary Figure 4**; nRF = 0.09), while Asteroid had 2,198 differences (**Supplementary Figure 5**) and achieved an nRF of 0.47. Therefore, as in simulation, Chrono-STA produced excellent supertrees from empirical datasets comprised of highly taxonomically restricted timetrees.

**Figure 7.**
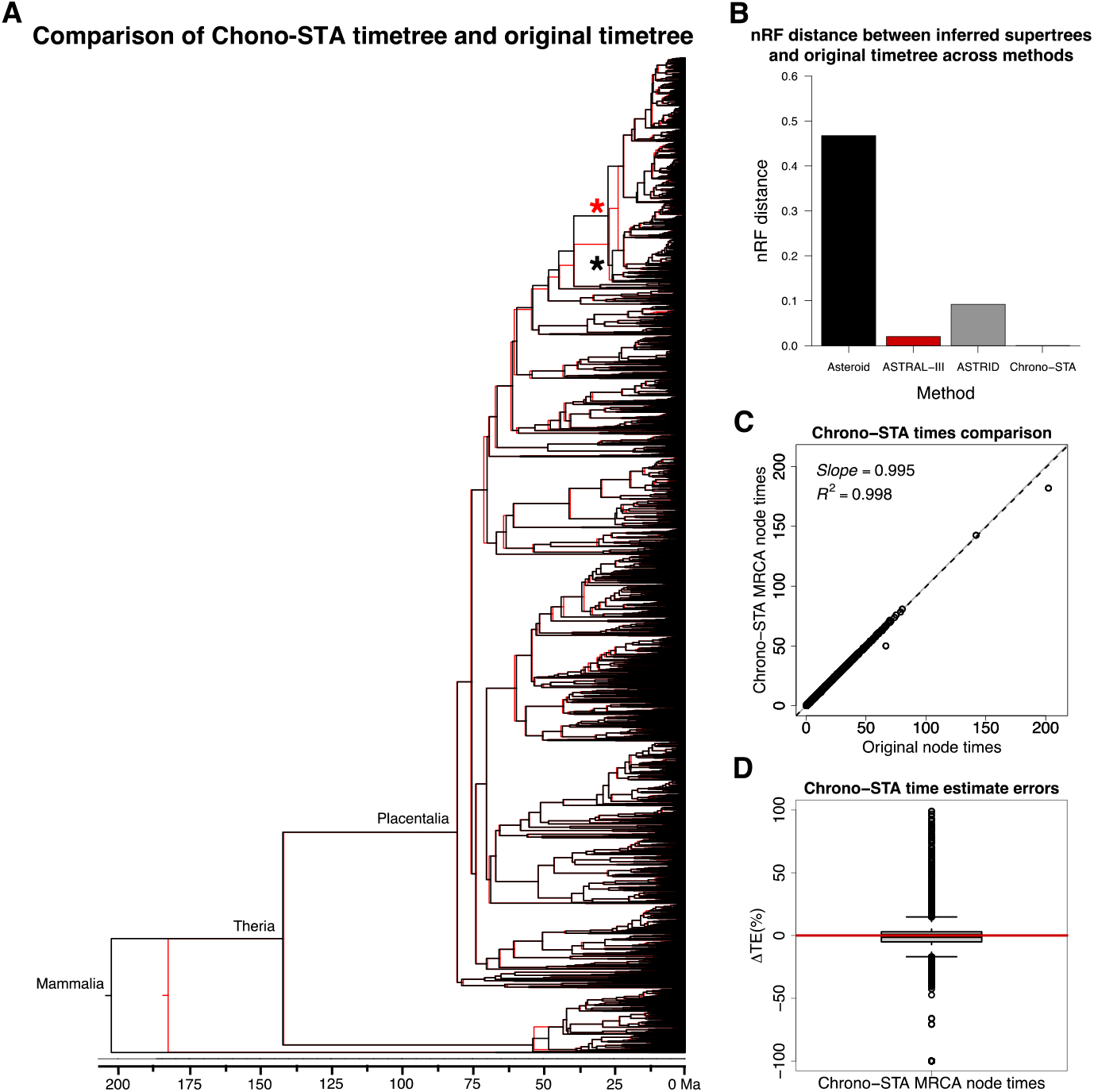
(**A**) Comparison of the 4,705 mammal species timetree (black) from Álvarez-Carretero et al. (2022) and Chrono-STA supertree (red). The supertree was constructed by combining 14 constituent timetrees including a backbone timetree. (**B**) nRF distances between the original timetree and the generated supertree for the 4,705 mammal species dataset (Álvarez-Carretero *et al*., 2022) for Asteroid (black), ASTRAL-III (red), ASTRID (grey), Chrono-STA (blue). The supertree was constructed by combining 14 constituent timetrees including a backbone timetree. The topological differences are marked with red and black asterisks. (**C**) Comparison of original and Chrono-STA time estimates. The *slope* and coefficient of determination (*R*^2^) for the linear regression through the origin are shown. The black dashed lines represent the best-fit linear regression through the origin. The solid gray line represents equality between estimates. (**D**) Boxplot of ΔTEs between the estimated and original node times. For Chrono-STA, we used the estimated node times for the MRCA of all the sets of taxa in the original timetree.

Chrono-STA recovered the correct node times as well, as they closely aligned with those of the original timetree, except for the Mammalia Chrono-STA node age (**Figure 7A**). This discrepancy arose because, in Chrono-STA, the calculation for that node time involved averaging across the 14 constituent timetrees, whereas in the original timetree, it represented the mammalian node time estimated independently for the backbone timetree. The *slope* and *R*^2^ values were nearly 1.0 (**Figure 7C**), and the median ΔTE was –0.29% (**Figure 7D**).

## 4 Discussion

We found that the new Chrono-STA approach can excel in cases where missing data are not randomly distributed among trees, but instead concentrated in certain clades (phylogenetically restricted). This better reflects the current state of the corpus of published literature, as researchers tend to specialize in certain families and genera and assemble detailed phylogenies of phylogenetically restricted groups. For such data with sparse species overlaps, the use of chronological information in times can help build better supertrees.

The performance improvement we observe from Chrono-STA as compared to the gene tree reconciliation approaches in building a supertree from phylogeny collections with phylogenetically restricted sparsity is likely due to the incorporation of time information. However, there was a large difference between the performance of ASTRAL-III and other methods (Asteroid and ASTRID). This difference likely arises from fundamental conceptual differences between them. ASTRAL-III (Zhang *et al*., 2018) combines phylogenies using a quartet-puzzling approach in which each constituent phylogeny is represented in batches of four taxa, and then the relative frequencies with which each of these quartets occur across all phylogenies are used to build the consensus supertree. In contrast, other Asteroid and ASTRID use distance between taxa in constituent phylogenies in the units of the number of intervening nodes or edges between taxa. When taxa are missing in some phylogeny, they impute missing distances statistically and then build a global matrix of pairwise distances to apply distance-based approaches, such as the Neighbor-Joining (Saitou and Nei, 1987), to construct a supertree.

Relative performance of many different versions of the imputation and quartet puzzling approaches have been examined for gene tree reconciliation with and without missing data (Liu and Warnow, 2023; Zhang and Mirarab, 2022; Rabiee *et al*., 2019; Zhang *et al*., 2020; Cao and Nakhleh, 2019). The general conclusion seems to be that they perform well and similarly. This is supported by our results, where ASTRAL-III consistently performed second-best after Chrono-STA, followed by a considerable gap in performance between ASTRAL-III and methods that used imputation to overcome missing data like Asteroid and ASTRID. This is not surprising because the reliability of any imputation is expected to be proportional to the amount of data available, resulting in more error when data are sparser. Furthermore, in cases of phylogenetically restricted sparsity, which, again, reflects the literature, this imputation is likely especially unreliable on a clade-by-clade basis, where there may be significantly less data than the matrix-wide average for some poorly studied clades. This would explain why ASTRAL-III, a quartet-puzzling approach that does not rely on imputation, achieves better accuracy than other methods except for Chrono-STA.

Therefore, Chrono-STA occupies a unique niche among supertree approaches, as it can reliably combine input constituent trees when the data are most sparse and when that sparsity is nonrandom. Compellingly, this condition is also likely the most reflective of the state of the literature across many large taxonomic groups, making Chrono-STA an attractive approach for reconstructing the history of life on Earth.

