## Supplementary Figures for "Integrating Phylogenies with Chronology to Assemble the Tree of Life"

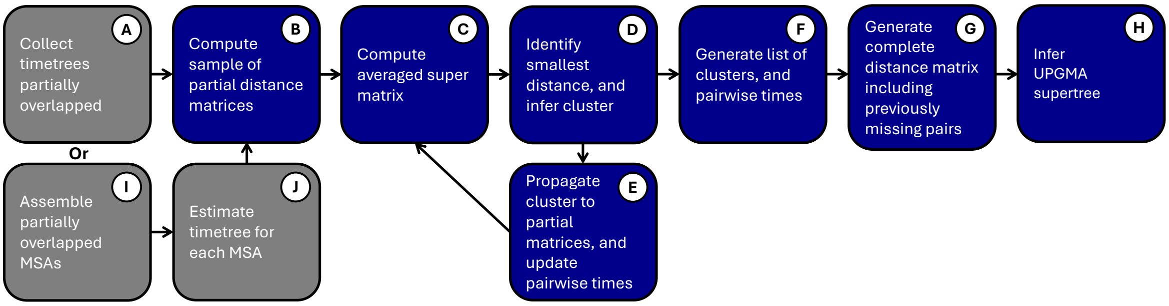


**Supplementary Figure 1.** Steps in the Chronological Supertree Algorithm (Chrono-STA). The process begins by collecting partially overlapped timetrees. A patristic pairwise time distance matrix is computed for each timetree. Then, an averaged supermatrix is computed. The smallest distance is identified, and a cluster is inferred. This cluster is then propagated to the partial matrices, updating pairwise times. The averaged supermatrix is recomputed, and the algorithm continues by iteratively clustering lineages until only one cluster remains. A list of clusters and pairwise distance times is generated. From this list, a complete pairwise time distances matrix, including previously missing pairs, is generated. Finally, a dated supertree is inferred by applying the unweighted pair group method with arithmetic mean (UPGMA). This approach can also be applied to combine trees from partially overlapped MSAs. In this instance, an ML tree is inferred for each MSA, followed by dating each ML tree. Then, the constituent timetrees are combined following the outlined procedure.

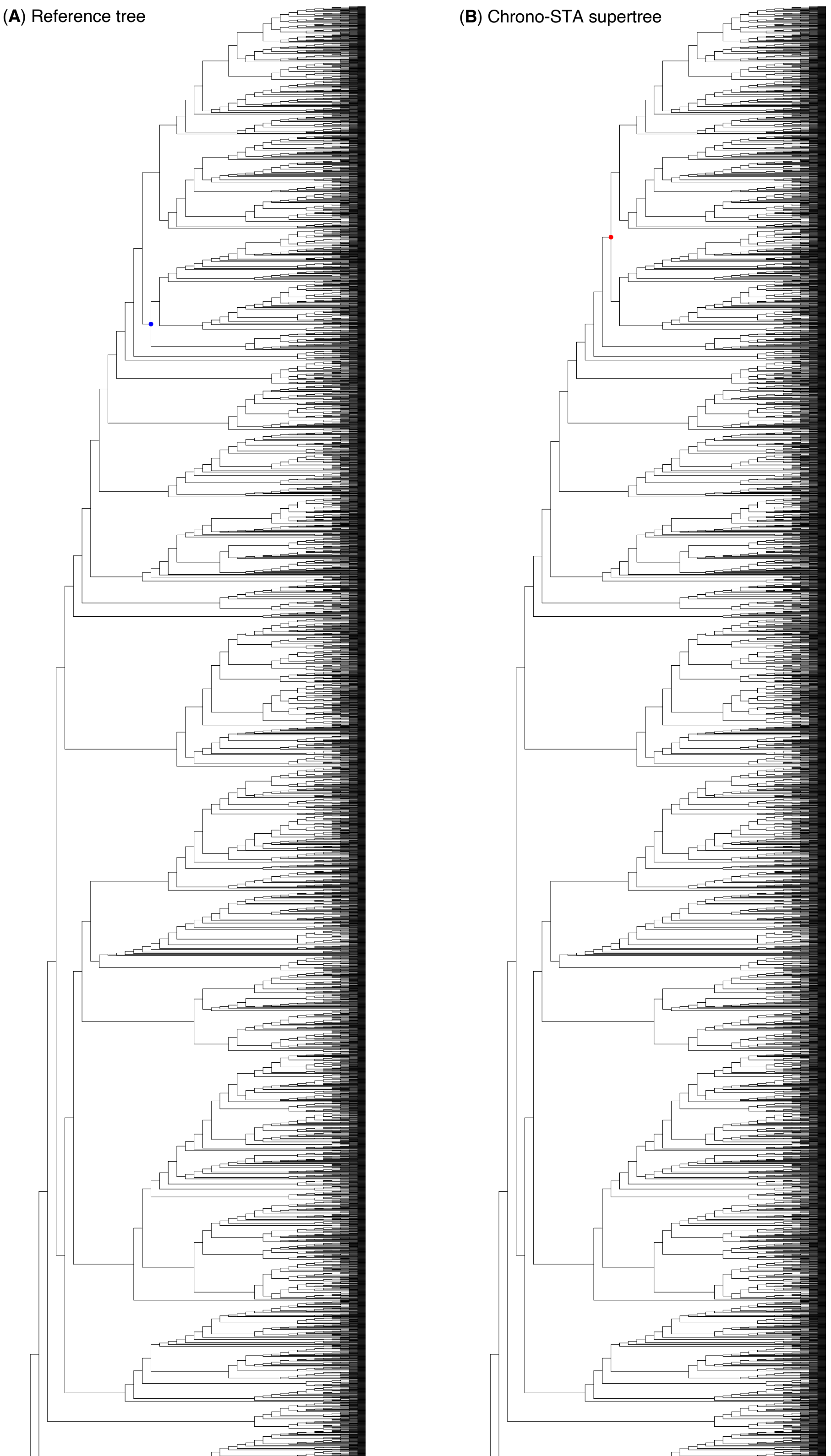


**Supplementary Figure 2.** Comparison of (**A**) the mammal timetree (Álvarez-Carretero *et al*., 2022) consisting of 4,705 species (reference tree) and (**B**) the Chrono-STA supertree. Red dots represent clades absent in (**A**), and blue dots represent clades absent in (**B**).

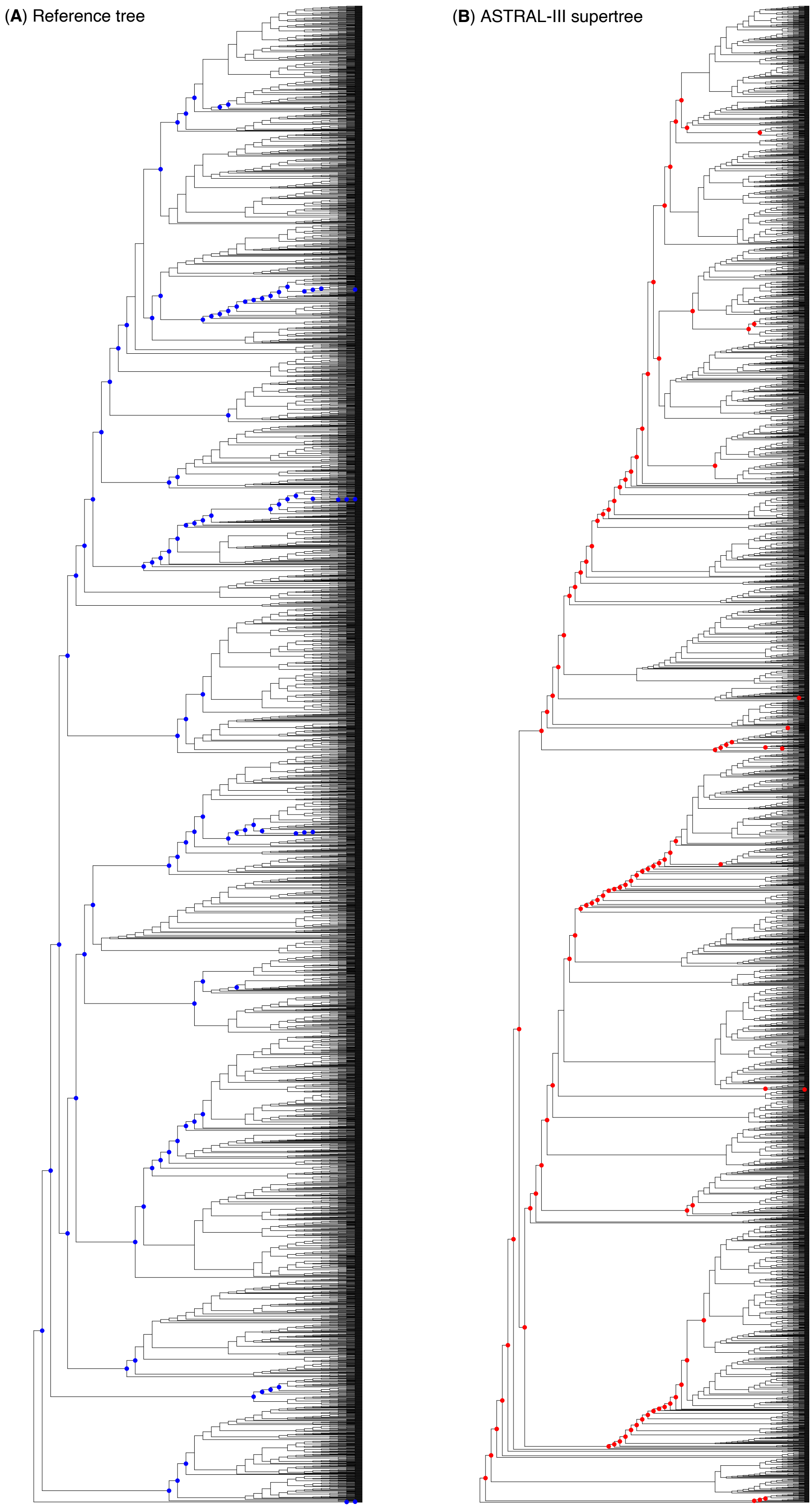


**Supplementary Figure 3.** Comparison of (**A**) the mammal timetree (Álvarez-Carretero *et al.*, 2022) consisting of 4,705 species (reference tree) and (**B**) the ASTRAL-III supertree. Red dots represent clades absent in (**A**), and blue dots represent clades absent in (**B**).

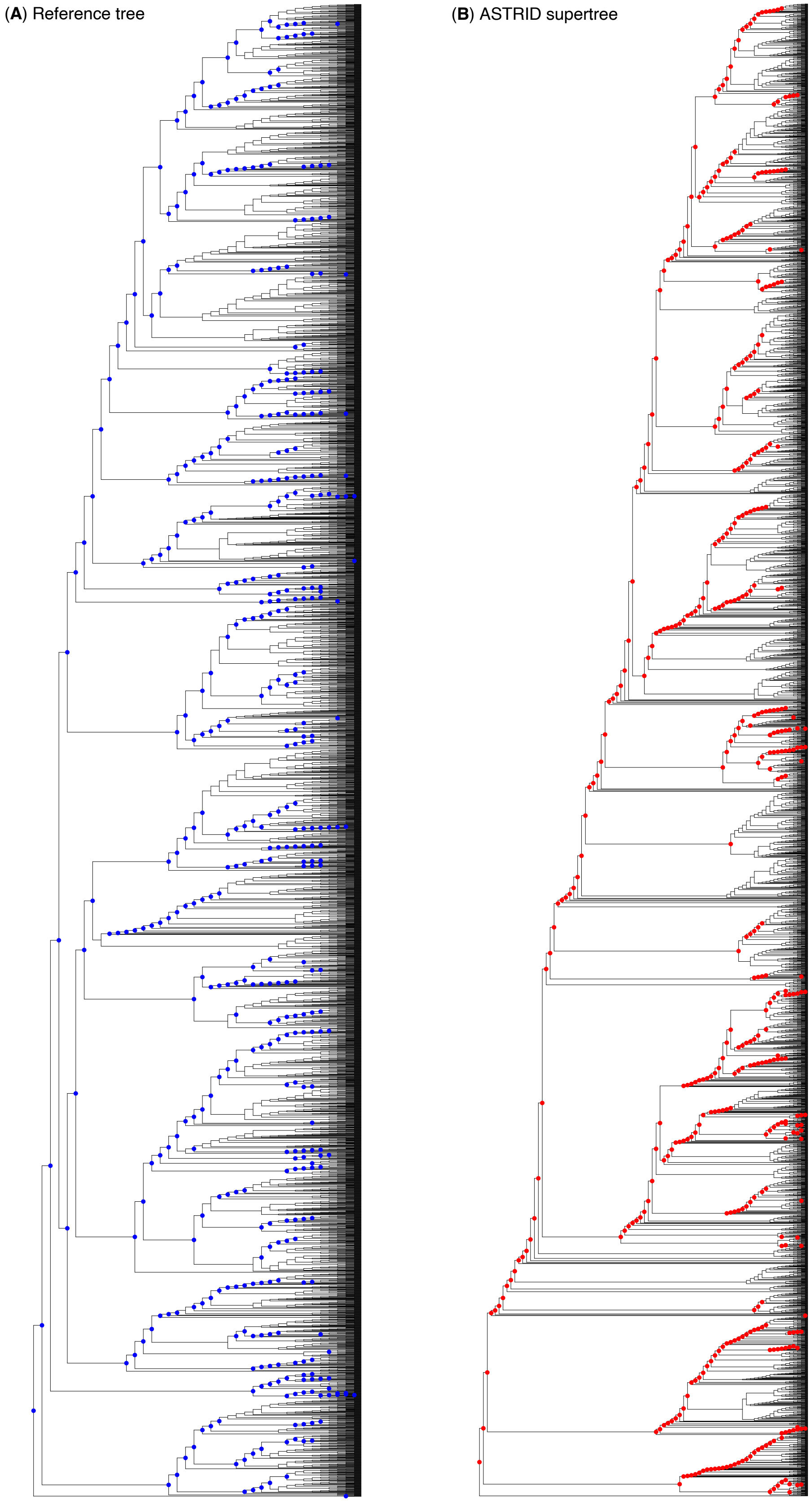


**Supplementary Figure 4.** Comparison of (**A**) the mammal timetree (Álvarez-Carretero *et al.*, 2022) consisting of 4,705 species (reference tree) and (**B**) the ASTRID supertree. Red dots represent clades absent in (**A**), and blue dots represent clades absent in (**B**).

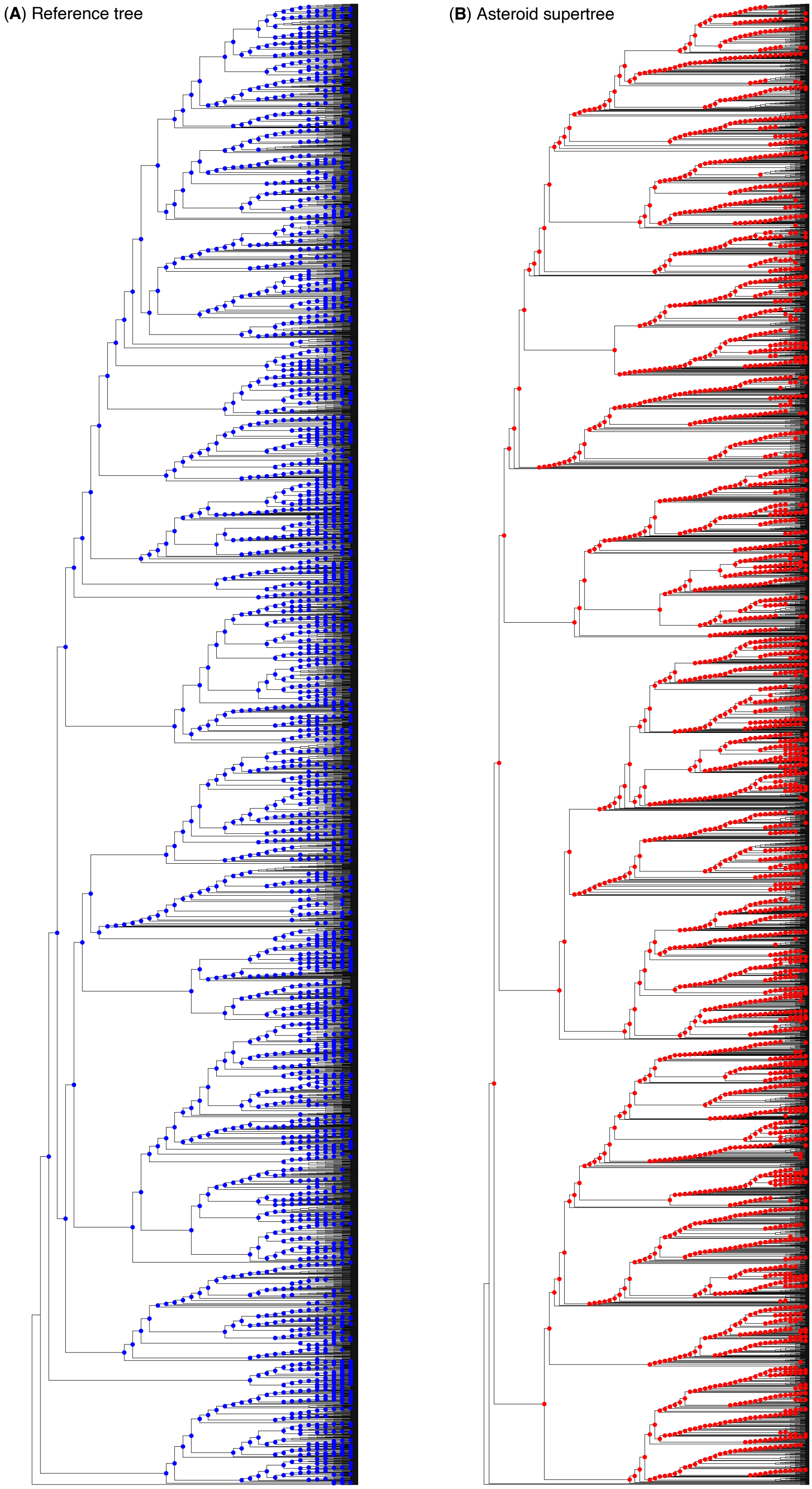


**Supplementary Figure 5.** Comparison of (**A**) the mammal timetree (Álvarez-Carretero *et al.*, 2022) consisting of 4,705 species (reference tree) and (**B**) the Asteroid supertree. Red dots represent clades absent in (**A**), and blue dots represent clades absent in (**B**).
